# Whole genome duplications and the diversification of the Globin-X genes of vertebrates

**DOI:** 10.1101/2021.03.28.437409

**Authors:** Federico G. Hoffmann, Jay F. Storz, Shigehiro Kuraku, Mike W. Vandewege, Juan C. Opazo

## Abstract

The globin superfamily of vertebrate genes is a textbook example of how the interplay between local gene duplications, whole-genome duplications, and regulatory changes can facilitate the evolution of novel protein functions. Almost every vertebrate examined possesses copies of hemoglobin and myoglobin in their genomes, and both cytoglobin and neuroglobin are present in the vast majority of vertebrate genomes surveyed as well. The phylogenetic distribution of globin-E, globin-Y and globin-X (GbX), however, is more spotty, suggesting multiple independent gene losses. Globin-X is an enigmatic globin with a wide phyletic distribution that spans protostomes and deuterostomes. Unlike canonical globins such as hemoglobins and myoglobins, functional data suggest that GbX does not have a primary respiratory function. Instead, available evidence suggests that GbX may play a role in protecting cells from oxidative damage and in reducing nitrite and it is predicted to be bound to the cell membrane. Recently released genomes from key vertebrate taxa provide an excellent opportunity to address questions about the early stages of evolution of these genes in vertebrates. In the current study, we integrate bioinformatic, synteny and phylogenetic analyses to characterize the diversity of GbX genes in non-teleost ray-finned fishes, resolve relationships between the GbX genes of cartilaginous fish and the GbX genes of bony vertebrates, and demonstrate that the GbX genes of cyclostomes and gnathostomes have independent duplicative histories. Our study highlights the role that whole genome duplications (WGDs) have played in expanding the repertoire of genes in vertebrate genomes. Our results indicate that GbX paralogs have a remarkably high rate of retention following WGDs in comparison to other globin genes, and also provide an evolutionary framework for interpreting results of experiments that examine functional properties of GbX and patterns of tissue-specific expression. By identifying GbX genes products of different WGDs in the vertebrate tree of life, our results can guide the design of experimental work to explore whether gene duplicates that originate via WGDs have evolved novel functional properties or expression profiles relative to singleton or tandemly duplicated copies of GbX.

## 1 Introduction

Globins are small, oxygen-binding hemoproteins found in all domains of life (Vinogradov et al., 2005). The globin superfamily of vertebrates provides an excellent example of how the interplay between local gene duplications, whole-genome duplications, amino acid and regulatory changes can fuel the evolution of novel protein functions (Storz et al., 2011, 2013; Keppner et al., 2020). The different types of globins in vertebrate genomes can be classified into four groups 1)-androglobin, 2)-neuroglobin,3) globin-X (GbX), and 4)-vertebrate-specific globins. The fourth category includes hemoglobin and myoglobin genes of gnathostomes, in addition to cytoglobin, globin-E, globin-Y, and the independently evolved hemoglobin and myoglobin genes of cyclostomes (Hoffmann et al., 2012a). Vertebrate-specific globins derive from a single ancestral gene present in the common ancestor of vertebrates, and their phylogenetic distribution among contemporary species reflects a complex history of lineage-specific duplications and deletions. Androglobin, the most recently discovered member of the vertebrate globins, is a chimeric protein that includes a rearranged globin domain that can be traced back to the common ancestor of choanoflagellates and animals (Hoogewijs et al., 2012). Neuroglobin also represents an ancient globin lineage that originated prior to the split between protostomes and deuterostomes (Burmester et al., 2000, 2002; Roesner et al., 2005; Dröge and Makałowski, 2011; Blank and Burmester, 2012; Hoffmann et al., 2012a), and in spite of intensive efforts, its physiological function remains a mystery (Fago et al., 2004; Ascenzi et al., 2014; Burmester and Hankeln, 2014; Keppner et al., 2020).

Almost every vertebrate examined possesses copies of hemoglobin and myoglobin in their genomes, and both cytoglobin and neuroglobin are present in the vast majority of vertebrate genomes surveyed as well (Hoffmann et al., 2011; Opazo et al., 2015). By contrast, the phylogenetic distribution of globin-E, globin-Y and GbX is more spotty, suggesting multiple independent gene losses. In the case of GbX, recently released genomes from key vertebrate taxa provide an excellent opportunity to address questions about its evolution. Globin-X is an especially enigmatic globin because it is predicted to be bound to the cell membrane (Blank et al., 2011), and because it has a very wide phyletic distribution that spans protostomes and deuterostomes (Blank and Burmester, 2012; Hoffmann et al., 2012a; Prothmann et al., 2020). There are GbX genes present in the genomes of insects, crustaceans, platyhelminthes, myriapods, spiders, hemichordates, and vertebrates among others, indicating that its origin predates the split between protostome and deuterostomes (Dröge and Makałowski, 2011). Unlike canonical globins such as hemoglobins and myoglobins, functional data suggest that GbX does not have a primary respiratory function. Instead, available evidence suggests that GbX may play a role in protecting cells from oxidative damage and in reducing nitrite (Corti et al., 2016; Koch and Burmester, 2016).

Whole-genome duplications have played a prominent role in the expansion and functional diversification of vertebrate-specific globins (Storz et al., 2011, 2013; Hoffmann et al., 2012b) and the hemoglobin gene repertoire of teleost fish (Opazo et al., 2013). Current evidence suggests that the repertoire of vertebrate GbX also expanded via WGDs. Initial genomic surveys of vertebrates revealed the presence of a single copy of GbX in a small number of distantly related vertebrate lineages that included some amphibians, some squamates, some teleost fish, elephant fish, and sea lamprey, which were all assumed to be 1-to-1 orthologs of each other (Roesner et al., 2005; Dröge and Makałowski, 2011; Hoffmann et al., 2012a). As the sample of vertebrate genomes increased, it became clear that there were different GbX genes in vertebrates (Opazo et al., 2015), and that apparent orthology among single copy GbX genes was the product of ‘hidden paralogy’, where genes are mistakenly identified as orthologs because of reciprocal, lineage-specific losses of alternative paralogs (Kuraku, 2010). Variation in the number of GbX paralogs among taxa and synteny comparisons suggest that whole-genome duplications (WGDs) were responsible for the expanded repertoire of GbX genes in vertebrates, and that subsequent lineage-specific WGDs also contributed to the increased GbX copy number in teleosts and salmonids (Opazo et al., 2015; Gallagher and Macqueen, 2017). However, questions remain regarding 1-) the diversity of GbX genes in non-teleost ray-finned fishes, 2-) the relationship of one of the GbX genes of cartilaginous fish and the GbX genes of bony vertebrates, and 3-) the relationships between the GbX genes of cyclostomes and gnathostomes. Accordingly, the goal of this study is to unravel the duplicative history and diversification of GbX in vertebrates to address these questions. In addition, we analyze newly released genomes from vertebrate groups that experienced additional rounds of WGD to track the evolutionary fate of the GbX genes. Specifically, we integrate synteny and phylogenetic analyses to decipher the evolution of the GbX repertoire of cyclostomes and cartilaginous fish, and we examine the role of WGDs in the diversification of the GbX gene repertoire in several fish and amphibian taxa that experienced lineage-specific WGDs subsequent to the two rounds of WGD in the stem lineage of vertebrates. Our phylogenies also identify a highly divergent GbX paralog in several teleost fish, which might reflect the emergence of a functionally distinct GbX protein.

## 2 Materials and Methods

### 2.1 Bioinformatic searches

We combined bioinformatic searches for GbX-like sequences in vertebrate genomes in the National Center for Biotechnology Information (NCBI) (Sharma et al., 2018) and the <u>Ensembl v.101 databases</u> (Yates et al., 2020), some of them coming from the Vertebrate Genomes Project (Rhie et al., 2020). Our searches were seeded with known GbX paralogs identified in Opazo et al. (2015) from coelacanth, elephant fish (*Callorhinchus milii*, which is also referred to as elephant shark), spotted gar, and zebrafish. We first retrieved all putative GbX orthologs and paralogs of vertebrates from <u>Ensembl v.101</u>. We then extended our searches to include additional vertebrate genomes available in NCBI from lineages not well-represented in previous studies (Hoffmann et al., 2012a; Opazo et al., 2015; Gallagher and Macqueen, 2017). Importantly, our surveys include a much wider array of vertebrate lineages, allowing us to perform a much more comprehensive survey of the diversity of their GbX repertoires. We now include multiple cyclostomes, multiple cartilaginous fish, multiple amphibians, more squamates, more teleost fish, recently released genomes from non-teleost ray-finned fish (Du et al., 2020; Bi et al., 2021), plus the tuatara (the single extant representative of the order Rhynchocephalia). Because WGDs appear to have played an important role in the expansion of the vertebrate GbX repertoire, we purposely included genomes from representatives of vertebrate groups that have undergone additional lineage-specific WGSs. Such taxa include the sterlet (Du et al., 2020), salmonids (Berthelot et al., 2014; Lien et al., 2016), members of the subfamily Cyprininae (Xu et al., 2014, 2019; Chen et al., 2019), and the African clawed frog (*Xenopus laevis*) (Session et al., 2016). In the case of the pacific lamprey, *Entosphenus tridentatus (Hess et al., 2020)*, the pouched lamprey, *Geotria australis*, and the southern lamprey, *Mordacia mordax*, we annotated GbX genes by pairwise comparisons with the GbX genes of the sea lamprey, *Petromyzon marinus*, using BLAST (Altschul et al., 1990) and the ‘Blast 2 sequences’ tool (Tatusova and Madden, 1999). Similarly, we used the GbX genes from elephant fish to search for unannotated GbX paralogs in additional genomes from other cartilaginous fishes. Finally, as outgroup sequences, we included the full repertoire of globins from the acorn worm (*Saccoglossus kowalevskii*, Hemichordata), an invertebrate representative of deuterostomes that possesses the most diverse globin repertoire in the group (Hoffmann et al., 2012a). We verified the identity of candidate GbX genes by reciprocal BLAST, comparing putative GbX sequences against the non-redundant protein sequence database (nr) of deuterostomes.

### 2.2 Sequence Alignment and Phylogenetic Analyses

We aligned amino acid sequences using the L-INS-i strategy from MAFFT v 7.471 (Katoh, 2005; Katoh et al., 2019) and estimated phylogenetic relationships using IQ-Tree v.2.0.6 (Minh et al., 2020). Support for the nodes was evaluated with the Shimodaira-Hasegawa approximate likelihood-ratio test and the aBayes tests (Anisimova et al., 2011) plus 10,000 pseudoreplicates of the ultrafast bootstrap procedure (Hoang et al., 2018). The best-fitting model of substitution was selected using the ModelFinder subroutine from IQ-Tree v.2.0.6 (Kalyaanamoorthy et al., 2017). Competing phylogenetic hypotheses were compared using the approximately unbiased test (Shimodaira, 2002a) as implemented in IQ-Tree v.2.0.6 (Minh et al., 2020).

### 2.3 Data curation

Because some of the sequences retrieved were annotated as neuroglobins or cytoglobins, and our searches potentially yielded redundant results, we first performed a phylogenetic analysis with all GbX candidates, to confirm the GbX identity of all retrieved sequences, to identify redundant sequences, and to detect potential annotation problems as evidenced by unusually long branches. Thus, for the second round of analyses, we removed all acorn worm globins other than 7, 8, 9,10, and 16, which had already been shown to be the most closely related to GbX (Hoffmann et al., 2012a; Prothmann et al., 2020). We removed redundant records and retained a single representative species per genus, with the exception of the carp, where we kept two separate assemblies that include two separate duplications. Finally, we also removed truncated genes from the dataset. In the cases of African clawed frog, medaka, and zebrafish, the Ensembl and NCBI sequences are almost identical, so we only kept one. In the cases of elephant fish, sea lamprey, and coelacanth, we removed the Ensembl sequences and we only used records derived from the NCBI database due to better coverage and availability of synteny information. For example, the tree in <u>supplementary figure S1</u> and synteny comparisons show that the truncated sea lamprey GbX paralog <u>ENSPMAG00000007241</u> from Ensembl, which comes from an earlier assembly (<u>Pmarinus_7.0</u>), corresponds to the full-length gene <u>LOC116943182</u> from NCBI, which comes from a more recent chromosome-level assembly (<u>kPetMar1.pri</u>) and includes better-resolved synteny. Finally, we discarded unusually short or long GbX candidates such as the <u>ENSMALG00000006192</u> gene from the swamp eel, *Monopterus albus*, which is 376 amino acids long, and all of the putative GbX genes from the southern lamprey. Details on data curation are provided in supplementary table S1.

### 2.4 Synteny analyses

We explored the genomic context of the GbX genes in the <u>Ensembl database v.100</u> (http://apr2020.archive.ensembl.org/index.html) by analyzing the presence of syntenic genes in vertebrate genomes with the help of the Genomicus browser v.100.01 (Nguyen et al., 2018). In the case of genomes not available in Ensembl, we checked synteny using the corresponding NCBI gene page, in combination with BLAST searches. Finally, because the presence of different PLEKHG paralogs has been used to define the genomic context of the different GbX paralogs of vertebrates (Opazo et al., 2015; Gallagher and Macqueen, 2017), we estimated phylogenetic relationships, following the same procedure mentioned above, for the PLEKHG1-3 paralogs of vertebrates to confirm our inferences of orthology. These analyses followed the same protocols used for the GbX phylogenies: we aligned amino acid sequences using the L-INS-i strategy from MAFFT v 7.471 (Katoh, 2005; Katoh et al., 2019) and estimated phylogenetic relationships using IQ-Tree v.2.0.6 (Minh et al., 2020) under the best-fitting model of substitution selected by the ModelFinder subroutine from IQ-Tree v.2.0.6 (Kalyaanamoorthy et al., 2017).

## 3 Results

### 3.1 Data description and nomenclature

We obtained 139 putative vertebrate GbX sequences from the <u>Ensembl</u> database, <u>release 101</u>, last accessed on September 28th, 2020, corresponding to the Ensembl gene tree <u>ENSGT00730000111686</u>, with representatives from jawless fish, ray-finned fish, squamates, testudines, tuatara, amphibians, and lobe-finned fish. We added an additional 25 sequences representing GbX candidates from testudines, amphibians, squamates, cartilaginous fish, lobe-finned fish, ray-finned fish, cyclostomes, plus the complete repertoire of globins from the acorn worm (Supplementary Table 1). As in previous studies, we did not find traces of GbX in the genomes of crocodilians, birds, or mammals despite the increased availability of genomes for these groups.

For the sake of consistency with previous studies, we followed the nomenclature from Gallagher and Macqueen (2017) in labeling orthologs of jawed vertebrates (gnathostomes). This nomenclature integrates information from phylogenetic and synteny analyses to infer orthology. Thus, orthologs of the spotted gar GbX1 gene <u>ENSLOCG00000014709</u>, which is flanked by PLEHG2 and SUPT5, were labeled as GbX1 genes, and orthologs of the spotted gar GbX2 gene <u>ENSLOCG00000012798</u>, which is flanked by PLEKHG3 and SRP14, were labeled as GbX2 genes. Within teleosts, which include duplicates of GbX2, orthologs of the GbX2a gene from Northern pike (*Esox lucius*, <u>ENSELUG00000004427</u>), which is flanked by a copy of PLEKHG3 and SRP14, were labeled as GbX2a genes, whereas orthologs of the GbX2b gene from the Northern pike (<u>ENSELUG00000016373</u>), which is flanked by PAPLNB and another copy of PLEKHG3, were labeled as GbX2b genes. Additional duplicates were identified by adding alternating letters and numbers to the name, as in the case of the sterlet GbX1a, GbX1b, GbX2a, and GbX2b genes, or the salmonid GbX2a1 and GbX2a2 genes. In these cases, similarities in the name of the paralogs do not reflect a shared duplicative history. In the case of cyclostomes, we labeled orthologs of the sea lamprey gene <u>LOC116943182</u> (flanked by PLEKHG3 and SRP14) as GbX-C2, and orthologs of the sea lamprey gene <u>LOC116948349</u> (flanked by another copy of PLEKHG3, TMEM180, and PYGL) as GbX-C1.

### 3.2 Phylogenetic analyses

We first estimated a maximum likelihood phylogenetic tree for the initial dataset of 190 sequences to ensure they were true GbX genes (available as Vert_GbX.190.fasta, Supplementary Online Material). The resulting tree placed all putative vertebrate GbX genes in a strongly supported monophyletic group that is sister to the clade that includes acorn worm globins 7, 8, 9,10, and 16, as in previous studies (Hoffmann et al., 2012a; Opazo et al., 2015), <u>Fig 1</u>, <u>supplementary figure 1</u>). This tree confirmed the GbX identity of all the sequences we retrieved, and placed the GbX1 genes of gnathostomes and the GbX-C1 and GbX-C2 genes of cyclostomes in monophyletic groups, but the gnathostome GbX2 sequences were paraphyletic relative to the GbX1, GbX-C1, and GbX-C2 genes. These initial analyses also identified a duplication of GbX1 in the sterlet, duplications of the GbX2 gene in the Leishan spiny toad and the sterlet, and duplications of the teleost GbX2a gene in the subfamily Cyprininae in addition to the duplication of the GbX2a paralog in salmonids that was previously identified by Gallagher and Macqueen (Gallagher and Macqueen, 2017). In the case of the Leishan spiny toad and the common carp, the two paralogs are found on the same genomic fragment (<u>Supplementary Table 1</u>), suggesting that they derive from single-gene tandem duplications. A closer inspection of the tree revealed several unusually long branches that may have been caused by the inclusion of low-quality sequences, such as the GbX paralogs from the southern lamprey, which were excluded from further analyses. Importantly, this analysis demonstrated that species within a genus share a common GbX repertoire. The only exception was the absence of a GbX2b paralog in the orange clownfish, *Amphiprion percula*, relative to the clown anemonefish, *Amphiprion ocellaris*. The estimated phylogeny in combination with synteny comparisons revealed the presence of redundant paralogs in coelacanth, elephant fish, Western clawed frog, sea lamprey, medaka, and zebrafish.

**Figure 1.**
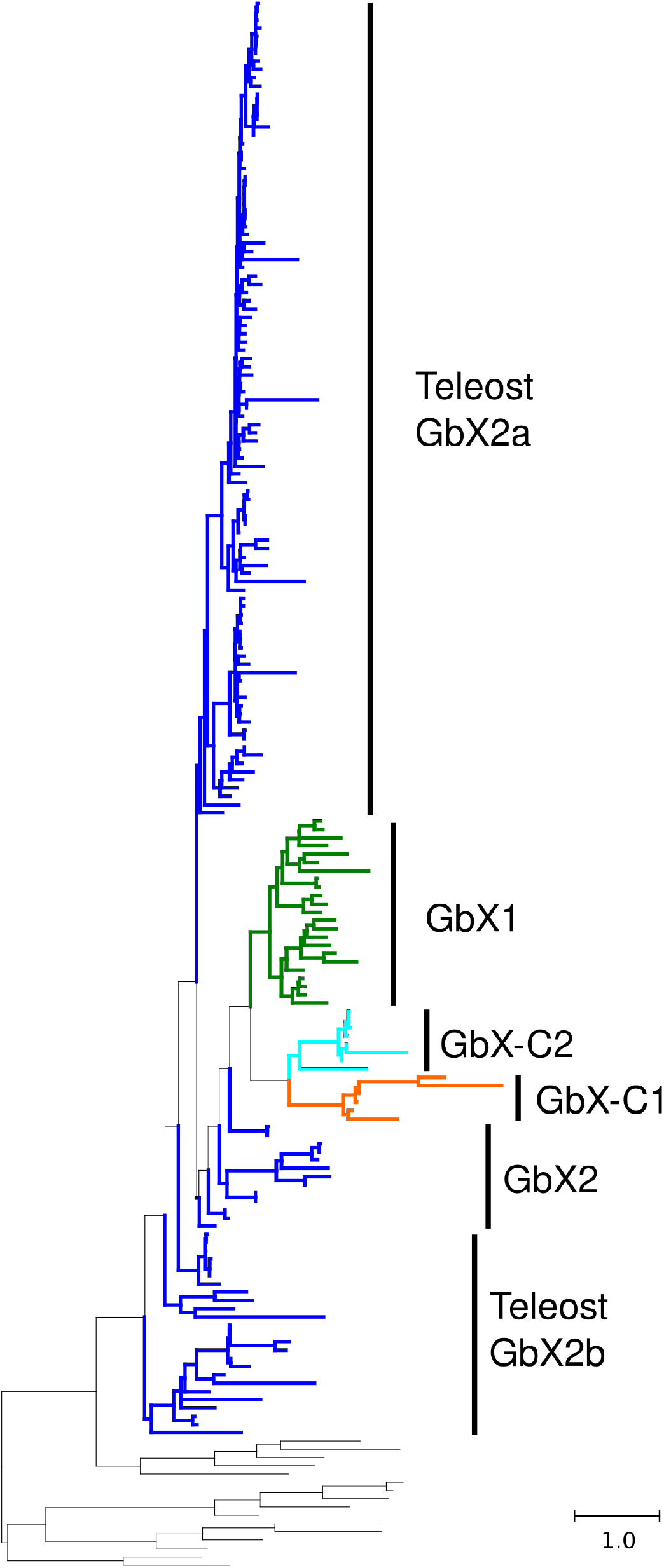
Maximum likelihood phylogram describing evolutionary relationships among the vertebrate globin-X candidates identified in our study. The tree was rooted with the full set of Acorn worm globins. The tree with the terminal labels is available as Supplementary Figure 1, and corresponding alignment is available as Supplementary Data File 1.

In the second round of analyses, we removed all acorn worm globins other than 7, 8, 9,10, and 16, we removed redundant records, we retained a single representative species per genus (with the exception of the carp, where we kept two separate assemblies that include two separate duplications), and we removed truncated genes (available as Vert_GbX.134.fasta, Supplementary Online Materia). As in the initial analysis, the resultant tree placed the GbX1 genes of gnathostomes and the GbX-C1 and GbX-C2 genes of cyclostomes in monophyletic groups, but GbX2 sequences were paraphyletic relative to the clade that included the GbX1, GbX-C1, and GbX-C2 genes, <u>supplementary figure 2</u>. In this tree, relationships for the GbX2 sequences deviated quite strongly from the expected organismal relationships. In particular, the GbX2a ohnolog (paralog derived from WGD) of teleost fish was split into three separate clades, and the GbX2b ohnolog of teleosts was split into multiple lineages that were placed as the deepest divergences of vertebrate GbXs. A strict reconciliation of this maximum likelihood tree with the organismal tree would imply the presence of over ten GbX paralogs in the last common ancestor of vertebrates with a large number of independent gene losses in descendent lineages, and would also imply that the duplication giving rise to the teleost GbX2a and 2b ohnologs traces back to the vertebrate ancestor. However, a tree that minimizes these independent deletions, where the GbX1 and GbX2 genes of gnathostomes were constrained to be monophyletic, and the GbX2a and GbX2b ohnologs of teleosts were constrained to be monophyletic within GbX2s and sister to each other, was not significantly different from the unconstrained tree (<u>Table 1</u>). Because this constrained tree minimizes the inferred number of independent gene gains and losses, it agrees well with assessments of conserved synteny among gnathostomes, and maps their duplication to the stem of the teleosts on the organismal tree, we selected it as most plausible phylogenetic hypothesis and use it as the basis of our evolutionary inferences. In this constrained tree (<u>Fig 2</u>, <u>supplementary figure 3</u>), the GbX1 and GbX2 paralogs of gnathostomes were placed sister to each other, and the GbX-C1 and GbX-C2 paralogs of cyclostomes were placed sister to each other as well.

**Table 1.** Results of topology tests

| Tree | logL | $\Delta L$ | bp-RELL | p-KH | p-SH | c-ELW | p-AU |
| --- | --- | --- | --- | --- | --- | --- | --- |
| Unconstrained | -20600.3 | 0 | 0.428 | 0.617 | 1 | 0.428 | 0.629 |
| GbX2 monophyletic | -20604.5 | 4.2 | 0.199 | 0.383 | 0.789 | 0.199 | 0.454 |
| GbX2, GbX2a, & GbX2b monophyletic | -20604.9 | 4.6 | 0.359 | 0.412 | 0.6 | 0.357 | 0.441 |
| GbX2 sister to GbX-C2 | -20640.4 | 40.1 | 0.0148 | 0.0466 | 0.121 | 0.015 | 0.036 |

bp-RELL : bootstrap proportion using REll method (Kishino et al., 1990).
p-KH : p-value of one sided Kishino-Hasegawa test (Kishino and Hasegawa, 1989).
p-SH : p-value of Shimodaira-Hasegawa test (Shimodaira and Hasegawa, 1999).
c-ELW : Expected Likelihood Weight (Strimmer and Rambaut, 2002)).
p-AU : p-value of approximately unbiased (AU) test (Shimodaira, 2002b).

**Figure 2.**
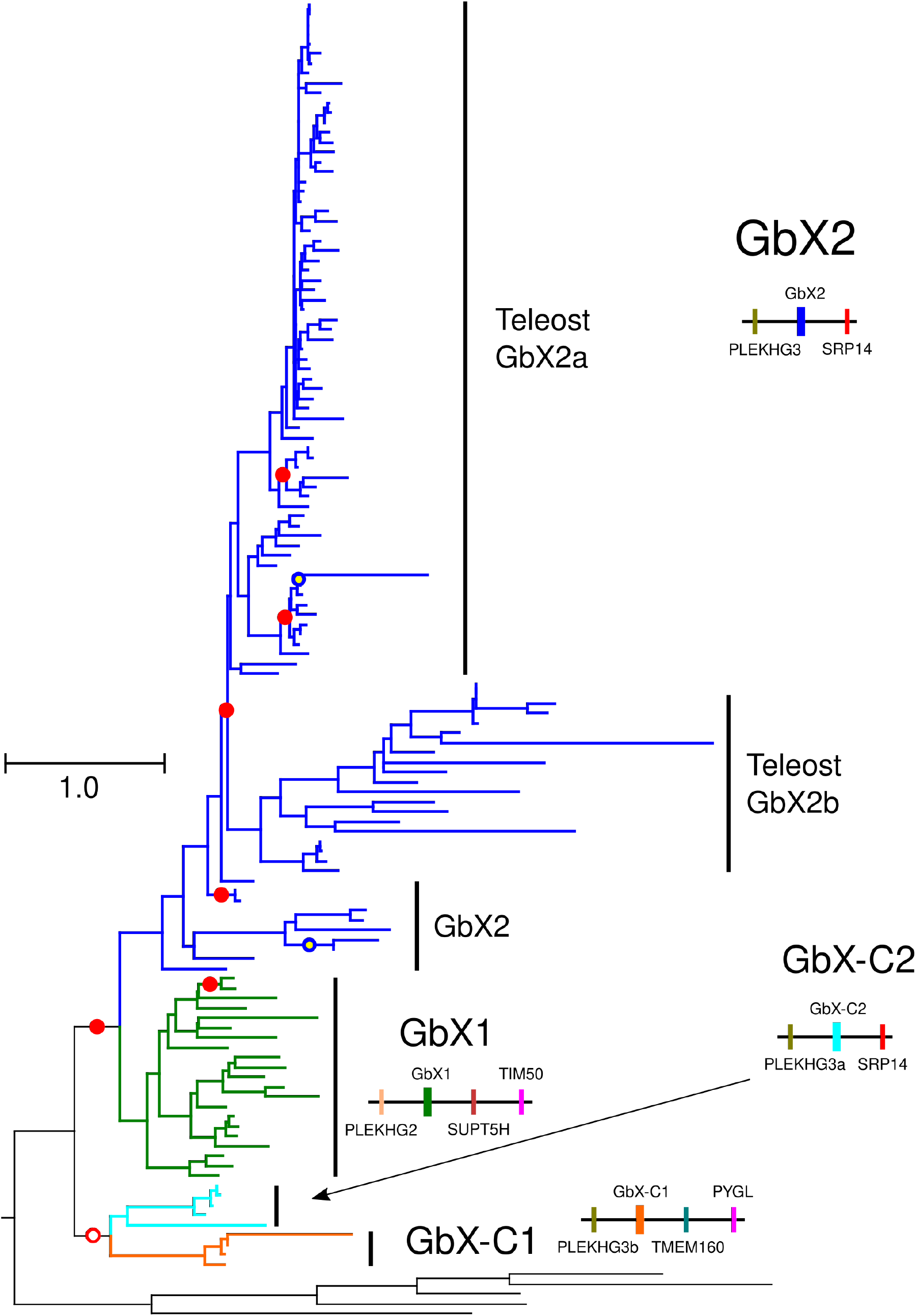
Maximum likelihood phylogram describing evolutionary relationships among the curated set of vertebrate globin-X candidates in our study, where the the GbX1 and GbX2 genes of gnathostomes were constrained to be monophyletic, and the GbX2a and GbX2b ohnologs of teleosts were constrained to be monophyletic within GbX2s and sister to each other. This tree was not statistically different from an unconstrained tree, which is available as Supplementary Figure 2, and minimizes the number of independent gene gains and losses. The tree was rooted with Acorn worm globins 7, 8, 9,10, and 16. The tree with the terminal labels is available as Supplementary Figure 3, and corresponding alignment is available as Supplementary Data File 2.

Each of the four lamprey species in the final analyses includes single-copy representatives of the GbX-C1 and GbX-C2 paralogs, and each of the paralog subtrees recovers the same species relationships. This pattern indicates that each of the cyclostome GbX paralogs can be traced back to the last common ancestor of these lamprey species. In addition, even though we only found a copy of GbX-C2 in hagfish, its position on the trees as sister to the clade of GbX-C2 sequences of lampreys, also supported by the available synteny data, indicates that the duplication that gave rise to GbX-C1 and C2 predates the split between hagfish and lampreys, and suggests that the hagfish secondarily lost the GbX-C1 paralog or that the apparent absence of the gene represents an assembly artifact.

In the case of gnathostome GbX genes, the GbX1 sequences from 1-cartilaginous fish, 2-ray-finned fishes, caecilians, 3-squamates, and 4-testudines were each placed in monophyletic groups; the single GbX1 gene from coelacanth is placed sister to the GbX1 genes from ray-finned fishes; and the GbX1 gene from tuatara is placed sister to the GbX1 genes from squamates (<u>supplementary figure 3</u>). As in previous studies (Opazo et al., 2015; Gallagher and Macqueen, 2017), we did not find traces of GbX1 in the genomes of teleost fishes in spite of our much denser sampling relative to earlier studies. The sterlet genome represents the only case where we identified duplicate copies of GbX1, and these were placed in a monophyletic group, sister to the GbX1 of the reedfish, with the spotted gar GbX1 as sister to them. Relationships among the GbX1 genes of ray-finned fishes were not congruent with known organismal relationships. The estimated gene tree placed the reedfish paralog sister to the 2 sterlet paralogs instead of spotted gar GbX1, but the corresponding branches were very short, so we ignored this discrepancy in our inferences. In this tree, caecilian GbX1 genes are placed sister to the clade that includes cartilaginous fish, ray-finned fish and lobe-finned fish sequences, instead of being placed sister to amniote GbX1s, but again, the corresponding branches were very short, so we did not attach importance to this apparent discrepancy. Relationships among the GbX1 sequences from amniotes matched the expected organismal relationships.

In the constrained tree (<u>Fig 2</u>, <u>supplementary figure 3</u>), relationships among the GbX2 genes matched organismal relationships at the order level. The putative elephant fish GbX2 ortholog is placed as sister to the clade that includes the GbX2 sequences from bony vertebrates, and relationships within the latter group were congruent with expected organismal relationships. The two tandem duplicates of the Leishan spiny toad are sister to each other in all analyses, and the two tandem duplicates of the carp are also close to each other in the tree, which suggests that they are very recent duplications (supplementary figures 1-3). The duplicate GbX2a paralogs of salmonids match organismal relationships (supplementary figures 1-3), consistent with the hypothesis that they trace their origins to the salmonid-specific WGD (Gallagher and Macqueen, 2017). The cyprinine-specific WGD appears to also have given rise to duplicate GbX2a paralogs in golden-line fish, carp, and goldfish. Here, however, the estimated tree groups the cyprinine paralogs by genus, with the exception of the tandem duplicate of the GbX2a paralog of the carp, <u>ENSCCRG00000022746</u>, which is grouped with the GbX2a paralogs of golden-line fishes (supplementary figures 1-3). A strict reconciliation of this subtree with the species tree would imply that golden-line fish (genus *Sinocyclocheilus*), goldfish (genus *Carassius*) and carp (genus *Cyprinus*) would each have undergone an independent WGD. However, our study involves a single gene family and we lack evidence for this from synteny comparisons.

Cyclostome genes are unusual with respect to nucleotide and amino acid composition and they also exhibit peculiar codon usage biases, which make it challenging to use phylogenetic approaches to resolve orthology between cyclostome and gnathostome genes (Qiu et al., 2011; Kuraku, 2013). In many instances, analyses of synteny have provided additional and independent information to resolve ambiguous gene phylogenies (Kuraku and Meyer, 2012; Campanini et al., 2015). In the case of GbX, the GbX2 genes of gnathostomes and the GbX-C2 genes of cyclostomes are flanked by copies of the PLEKHG3 and SRP14 genes, strongly suggesting they are 1-to-1 orthologs. However, a phylogenetic scenario where gnathostome GbX2 was constrained to be sister to the cyclostome GbX-C2 (as would be expected if the two genes were 1-to1 orthologs) was statistically rejected in our tree topology tests (Table 1).

Synteny analyses in Opazo et al. (2015) suggest that the GbX1 and GbX2 genes of gnathostomes derived from one of the two rounds of WGD early in vertebrate evolution. If the GbX2 of gnathostomes and the GbX-C2 of cyclostomes derived from the same duplication, the syntenic genes that co-duplicated with them would be expected to reflect the same duplicative history, so the PLEKHG3 gene of gnathostomes that is adjacent to GbX2 would be sister to the PLEKHG3 gene of cyclostomes that is adjacent to GbX-C2. We explored this issue by estimating phylogenetic relationships among the gnathostome and cyclostome PLEKHG2 and PLEKHG3 genes, which are used to define the genomic context of the vertebrate GbX genes and validate inferences of orthology (Opazo et al., 2015; Gallagher and Macqueen, 2017). The estimated PLEKH phylogeny recapitulates the results of the GbX phylogeny, placing the two PLEKHG3 paralogs of cyclostomes in a monophyletic group, as their neighboring GbX-C1 and GbX-C2 genes in the GbX tree, and placing the PLEKHG2 and PLEKHG3 genes of gnathostomes in a monophyletic group as their neighboring GbX1 and GbX2 genes in the GbX phylogeny. Thus, both synteny and phylogenetic analyses indicate that the GbX-C1 and C2 genes of cyclostomes and the GbX1 and GbX2 genes of gnathostomes are products of independent duplication events.

### 3.3 Expression of the different GbX paralogs

The available evidence indicates that the GbX paralogs of elephant fish, spotted gar, and salmon are differentially expressed (Opazo et al., 2015; Gallagher and Macqueen, 2017). In the elephant fish, RNA-seq data indicates that the GbX1 and GbX2 paralogs are mostly expressed in the spleen, and GbX1 (AKU74647), which is labeled as GbX2 in the original study of Opazo et al. (2015), is also expressed in brain, spleen and testis, whereas GbX2 (XP 007891388), which is labeled a GbX1 in Opazo et al. (2015), is expressed in brain, gills, intestine, kidney and liver (Opazo et al., 2015). In the case of spotted gar, quantitative PCR data indicate that GbX1 is most highly expressed in the brain, and expression is also detected in heart, gill, liver, intestine and spleen, whereas expression of GbX2 is only detected in the brain (Gallagher and Macqueen, 2017). In the case of salmonids, quantitative PCR data revealed the following: 1) expression of the GbX2a1 paralog (<u>ENSSSAG00000003360</u>) is highest in the brain and it is also expressed in intestine and eye; 2) expression of the GbX2a2 paralog (<u>ENSSSAG00000007165</u>) was not detected; and 3) expression of the GbX2b paralog (<u>ENSSSAG00000047904</u>)is highest in the intestine and is also detected in the brain, stomach, and eye (Gallagher and Macqueen, 2017). These data suggest that these genes are expressed in a variety of tissues, but that patterns of expression vary in a lineage-specific manner that is not easy to put into the evolutionary framework of the organismal tree at this stage.

## 4. Discussion

After performing an exhaustive homolog search using genome-wide sequence resources, we integrated phylogenetic and synteny analyses to infer the duplicative history of GbX paralogs in vertebrates. Because these analyses included highly contiguous cyclostome genomes plus newly released genomes from cartilaginous fish as well as representatives of the deepest-branching lineages of ray-finned fishes (Du et al., 2020; Bi et al., 2021), we were able to resolve unanswered questions regarding the early stages of evolution of GbX genes in vertebrates. Since the time of its initial discovery (Roesner et al., 2005), GbX went from being an obscure gene found in a very limited sample of vertebrates to becoming a credible candidate to provide clues about the functional role of the ancestor of all animal globins (Blank et al., 2011; Song et al., 2020). This change in paradigm came with an increased interest in deciphering its evolutionary history and its still elusive functional role (Burmester and Hankeln, 2014; Keppner et al., 2020). We have recently documented the presence of GbX paralogs in arthropods (Prothmann et al., 2020), confirming phylogenetic predictions that indicate that the origin of GbX predates the split between deuterostomes and protostomes (Burmester et al., 2002; Roesner et al., 2005; Dröge and Makałowski, 2011; Blank and Burmester, 2012; Hoffmann et al., 2012a)(Opazo et al., 2015), which is estimated to have occurred ~ 730 million years ago (Kumar et al., 2017).

### 4.1 Evolution of GbX in early vertebrates

Our reconstructions shed light on the early stages of evolution of the GbX genes in vertebrates. Our increased sampling allows us to resolve orthology for the two different GbX genes of cartilaginous fish relative to the rest of the gnathostomes. In turn, this implies that the two GbX paralogs of gnathostomes, GbX1 and GbX2, can be traced back to the last common ancestor of cartilaginous fish and bony vertebrates, and synteny analyses suggest that these two paralogs are actually ohnologs that derive from one of the two possible rounds of WGD early in vertebrate evolution, in agreement with Opazo et al. (2015). Similarly, the two GbX paralogs of cyclostomes, GbX-C1 and GbX-C2, can be traced back to the last common ancestor of hagfishes and lampreys. Synteny analyses of the GbX genes in the sea lamprey and the pouched lamprey reveal the shared presence of PLEKHG3 genes next to GbX-C1 and GbX-C2. The conserved synteny and phylogenetic analyses suggest that the PLEKHG3 and GbX genes of cyclostomes coduplicated, which would indicate that they derive from a segmental duplication or even a WGD. In this regard, our results are consistent with previous studies that indicate that lamprey genomes underwent extensive segmental duplications (Smith and Keinath, 2015), or even an additional and independent WGD (Mehta et al., 2013), although other scenarios are also possible.

Our results provide strong evidence that the gnathostome GbX paralogs derive from one of the two vertebrate-specific WGDs, either 1R or 2R, confirming inferences from Opazo et al. (2015) (<u>Fig 3</u>). In addition, the GbX-C1 and C2 paralogs appear to derive from segmental duplications involving additional genes, and this segmental duplication could correspond to a WGD. Reconciling the observed relationships between GbX1, GbX2, GbX-C1 and GbX-C2, with the organismal phylogeny is not trivial, especially given uncertainty about the timing and number of WGDs that occurred early in vertebrate evolution. There is agreement that early vertebrates underwent 2 rounds of WGD (Meyer and Schartl, 1999; McLysaght et al., 2002; Dehal and Boore, 2005), 1R and 2R, and there is also agreement that cyclostomes and gnathostomes share 1R. There is less agreement about the placement of 2R on the vertebrate tree (Kuraku et al., 2009). Some authors place 2R in the common ancestor of cyclostomes and gnathostomes (Sacerdot et al., 2018), whereas others place 2R in the last common ancestor of gnathostomes (Simakov et al., 2020), and there is also the suggestion that cyclostomes might have undergone an additional and independent WGD early in their evolution (Mehta et al., 2013). Our results do not fit any of these scenarios better than the alternatives, and would require multiple gene losses to fit in any of them.

**Figure 3.**
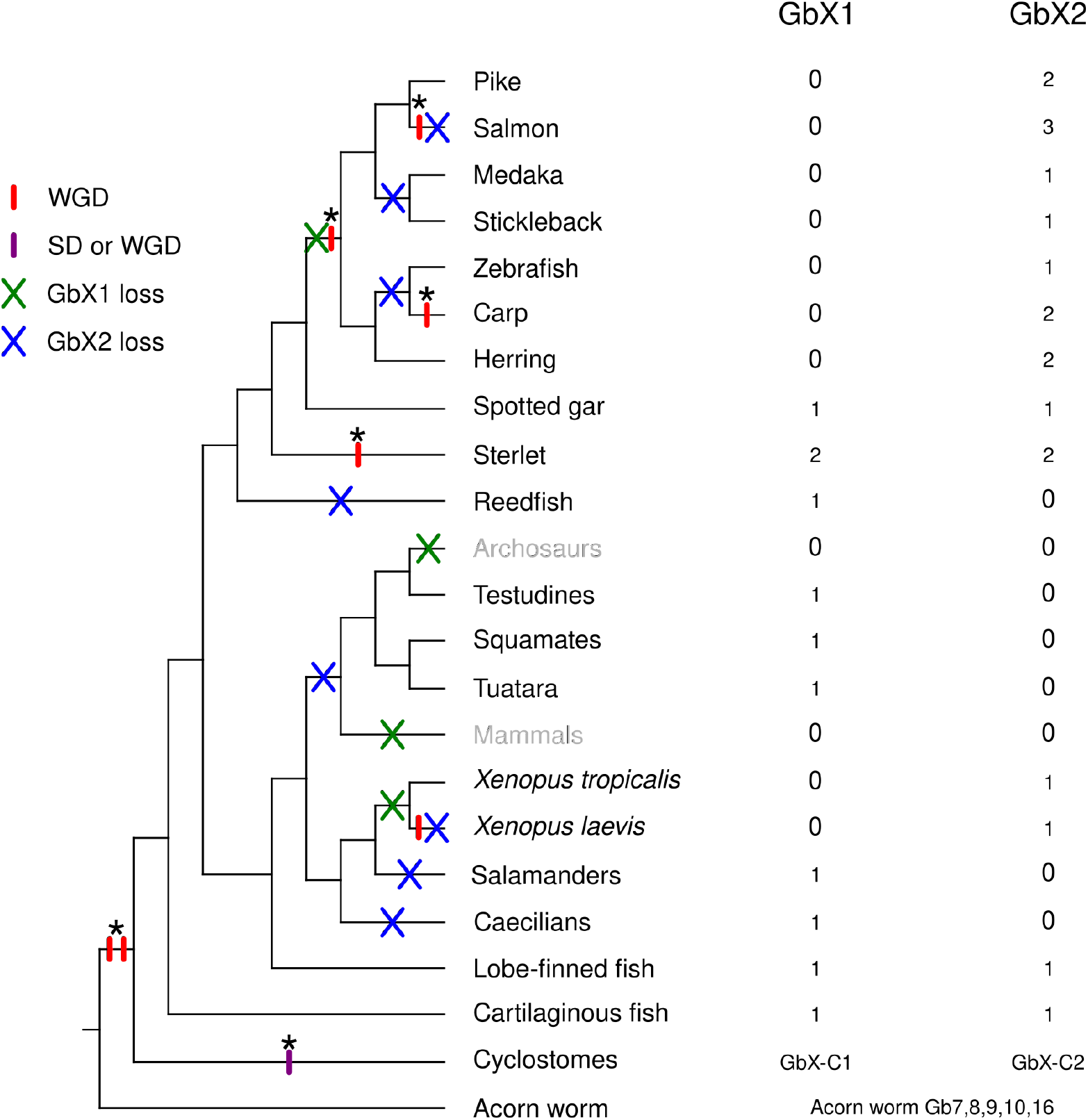
Graphical summary of the role of WGDs in the expansion of the vertebrate GbX repertoire. Organismal relationships on the right, and the number of GbX paralogs per lineage on the left. WGDs, segmental duplications (SD), and gene losses are mapped to their corresponding branch. We placed the 1R and 2R WGDs following Sacerdot et al. (2018). Symbols on a branch are arranged according to their relative order. Asterisks on top of the vertical bars denote that these duplications gave rise to GbX paralogs present in extant species. Because we cannot determine which of the 1R or 2R WGD in the vertebrate ancestor gave rise to GbX1 and GbX2,we placed the asterisk between them. derive from either 1R or 2R, and our analyses cannot determine which of the two. In the case of cyclostomes, our analyses indicate that GbX-C1 and GbX-C2 probably derive from a segmental duplication or a WGD. Note that a full tree of all species examined would include 2 additional tandem duplications and multiple additional gene losses.

### 4.2 Evolution of the GbX paralogs of gnathostomes and cyclostomes

The GbX paralogs of gnathostomes and cyclostomes have followed contrasting evolutionary trajectories. Both the GbX1 and GbX2 paralogs of gnathostomes and the GbX-C1 and GbX-C2 paralogs of cyclostomes can be traced back to the common ancestor of each group, but whereas the GbX1 and GbX2 paralogs of gnathostomes have been lost independently multiple times and have undergone additional duplications, the GbX-C1 and GbX-C2 paralogs of cyclostomes have been retained by all lampreys (<u>Fig 3</u>). It appears that hagfishes may have secondarily lost GbX-C1, but a more contiguous assembly is needed for confirmation. Among gnathostomes, spotted gar, elephant fish, sterlet, and coelacanth have retained both GbX1 and GbX2 paralogs, whereas mammals and archosaurs (birds + crocodilians) have lost both. GbX1 was independently lost in teleost fish, mammals, anurans, and archosaurs, whereas GbX2 was independently lost in caecilians, amniotes, and reedfish. Transcriptomic data indicate that only GbX1 has been retained in salamanders (Queiroz et al., 2020), which would imply an additional independent loss of GbX2.

Among ray-finned fish, the sterlet possesses a total of 4 GbX copies - the most of any vertebrate examined to date - as this species has retained both of the GbX1 and GbX2 duplicates derived from the WGD specific to this lineage (Du et al., 2020). Reedfish has only retained a copy of GbX1, gar has retained copies of both GbX1 and GbX2, and teleosts have only retained copies of GbX2 (<u>Fig 1</u>, supplementary figures 1-3). The GbX2a and GbX2b paralogs of teleost fish appear to derive from the teleost-specific WGD (Gallagher and Macqueen, 2017), and they have also been differentially retained among species. The GbX2 paralog has been retained in 83 out of 84 teleost fish in ensembl v101. The only exception is the Chinese medaka (*Oryzias sinensis*), and because of the overall low gene coverage of this assembly, we suspect this to be an artefact. By contrast, the GbX2b paralog has been retained in 25 out of 84 teleost fish genomes examined — representing 8 different higher-level lineages and implying multiple independent losses (Supplementary Table S1). The GbX2a and GbX2b paralogs of salmonid fishes also exhibit a highly asymmetric rates of gene retention: all six examined salmonid genomes retained the two GbX2a ohnologs derived from the salmonid-specific WGD (Gallagher and Macqueen, 2017), but only a single copy of GbX2b, suggesting that the latter gene reverted to a diploid state shortly after the salmonid-specific WGD (supplementary figures 1-3). Similarly, the duplicate GbX2 paralogs of the subfamily Cyprininae had contrasting fates. The GbX2a paralog duplicated in the cyprinine-specific WGD and has been retained in most cases, whereas the GbX2b paralog was apparently lost earlier, in the common ancestor of cyprinines and zebrafish.

In addition to showing contrasting levels of synteny conservation and retention rates, the GbX2a and GbX2b paralogs of teleost fish also seem to follow diverging evolutionary pathways in terms of genomic context and amino acid changes. The genomic context of the teleost fish GbX2a gene is highly conserved, and in the vast majority of the cases, the GbX2a gene is flanked by SRP14 and PLEKHG3, as in the ancestral GbX2 gene. In the case of GbX2b, however, the genomic context is more variable, although in most cases there is a copy of the CEP170 gene in the vicinity, and sometimes there is a copy PLEKHG3 as well. The GbX2a and GbX2b genes also exhibit different rates of amino acid substitution rates, as evidenced by the longer branches in the GbX2b portion of the phylogenetic trees. Asymmetric rates of evolution are often associated with differences in evolutionary constraints among sister paralogs, and the GbX2a and 2b paralogs of teleost fish appear to fit this pattern well (Pál et al., 2006).

Our synteny and phylogenetic analyses indicate that WGDs have played a major role in the diversification of GbX paralogs, just as they did in the vertebrate-specific globins (Hoffmann et al., 2012b; Opazo et al., 2013, 2015; Storz et al., 2013). Also like in vertebrate-specific globins, the repertoire of GbX in extant lineages has been impacted by lineage-specific losses and the differential retention of relatively old duplicates. Specifically, our results indicate that WGDs have given rise to at least 6 pairs of duplicates, that GbX1 was lost at least 4 independent times, and that GbX2 was lost at least 8 times independently (<u>Fig 3</u>), plus multiple additional losses of the GbX2b gene in teleost fishes. Starting with the oldest, the first pair of WGD-derived duplicates corresponds to the GbX1 and GbX2 ohnologs of gnathostomes, which derive from either the 1R or 2R WGDs early in vertebrate evolution. The second pair corresponds to the GbX2a and GbX2b ohnologs of teleost fish, which derive from the teleost WGD, and the third corresponds to the duplicate copies of GbX2a in salmonids, which derive from the salmonid-specific WGD, both identified by Gallagher and Macqueen (Gallagher and Macqueen, 2017). The fourth pair corresponds to the duplicate copies of GbX2a in cyprinins, which derive from the cyprinin-specific WGD, and finally, the fifth and sixth pair correspond to duplicate copies of both GbX1 and GbX2 that derive from the sterlet-specific WGD. The *Xenopus*-specific WGD is the only vertebrate WGD that does not appear to have produced an expansion of the GbX gene repertoire, and the possibility that the GbX-C1 and GbX-C2 duplicates of cyclostomes derive from an WGD as well.

Our study highlights the role that WGDs have played in expanding the repertoire of genes in vertebrate genomes. Our results indicate that GbX paralogs have a remarkably high rate of retention following WGDs in comparison to other globin genes. Our results also provide an evolutionary framework for interpreting results of experiments that examine functional properties of GbX and patterns of tissue-specific expression. By identifying GbX ohnologs that are products of different WGDs during the radiation of vertebrates, our results can guide the design of experimental work to explore whether gene duplicates that originate via WGDs have evolved novel functional properties or expression profiles relative to singleton or tandemly duplicated copies of GbX in the same species.

## 6 Author contributions

FGH designed the study. FGH, SK, JCO, and MWW collected and/or analyzed data. FGH, SK, JCO, and JFS wrote the manuscript. FGH, SK, JCO, and JFS reviewed and edited the manuscript. All authors contributed to the article and approved the submitted version.

## 8 Acknowledgements

FGH thanks Amanda Coward Black for editorial assistance. JCO wants to acknowledge the members of the Integrative Biology Group, Universidad Austral de Chile for their constant support, scientific enthusiasm, and creative feedback.

---

Supplementary Figure 1. Unconstrained Tree with 190 sequences with labels

Supplementary Figure 2. Unconstrained Tree with 134 sequences with labels

Supplementary Figure 3. Constrained Tree with 134 sequences with labels

Supplementary Figure 4. PLEKHG tree. We might need to move this figure to the main text.

